# Differential Impact of Isoflurane and Propofol on Apoptotic Regulation of Helper T cells

**DOI:** 10.64898/2026.08.18.745422

**Authors:** Priyanka Saha, Deepa Chakrabarti, Deepanwita Das, Meghna Mukherjee, Srabasti Barai, Sukanya Ghosh, Anurima Samanta, Dona Sinha

**Affiliations:** Dept. of Receptor Biology and Tumor Metastasis, Chittaranjan National Cancer Institute, 37, S.P. Mukherjee Road, Kolkata 700026; Dept. of Anaesthesiology, Chittaranjan National Cancer Institute, Street Number 299, DJ Block, Action Area I, Newtown, Kolkata 700156; Dept. of Epidemiology and Biostatistics, Chittaranjan National Cancer Institute, 37, S.P. Mukherjee Road, Kolkata, India

**Keywords:** Isoflurane, Propofol, CD4□ Th cells, Jurkat T cells, Apoptosis

## Abstract

**Background:** Anesthetic agents administered during surgery are one of the key perioperative factors affecting immune modulation in cancer patients. This comparative study elucidated the mechanisms by which the volatile anesthetic, isoflurane and the intravenous anesthetic, propofol impacted apoptosis signaling in CD4^+^ helper T (Th) cells.

**Methods:** Flow cytometry was used to analyze apoptosis, mitochondrial function and reactive oxygen species (ROS) generation, while Western blotting, ELISA and RT-qPCR were employed to study protein/gene expression in sorted CD4□ Th cells from perioperative breast cancer female patients (anesthetized with isoflurane or propofol, n=15 per group) and Jurkat T cells.

**Results:** Patient-derived CD4□ Th cells and Jurkat T cells exhibited that isoflurane at clinically relevant concentrations triggered apoptosis through mitochondrial depolarization, ROS generation, DNA damage, and activation of caspase-3/7. Specific use of caspase-3/7 inhibitor, Z-DEVD-FMK and antioxidant N-acetyl cysteine rescued isoflurane-induced apoptosis. Further, isoflurane relative to propofol, activated p38 mitogen-activated protein kinase (MAPK), and use of p38 inhibitor, SB203580 suppressed isoflurane-induced apoptosis. Collectively, these findings validated the involvement of the ROS-p38-caspase-3/7 axis in isoflurane-associated apoptosis signaling. On the other hand, propofol conserved mitochondrial integrity, reduced oxidative stress, and maintained higher proliferative capacity. Interestingly, isoflurane-associated apoptosis was transient, with postoperative recovery in patients and similar rescue from apoptosis was evident in Jurkat T cells within 24-48 h of drug removal.

**Conclusions:** By integrating analyses of patient-derived CD4^+^ Th cells with mechanistic validations in Jurkat T cells, this study identified the ROS-p38-caspase-3/7 signaling axis and the reversible nature of isoflurane-induced apoptosis.

## Introduction

The perioperative period represents a crucial window during which surgical stress and anaesthetic agents may influence host immunity and tumor progression. Intravenous anaesthetic, propofol has been associated with increased infiltration of T lymphocytes and natural killer cells [1] in breast cancer and with greater T helper (Th) cell activation and Th1 cell differentiation relative to volatile anaesthetic, isoflurane in non-small-cell lung cancer patients [2]. Perioperative immunosuppression may facilitate tumor cell dissemination and micrometastasis, thereby increasing the risk of recurrence or poor prognosis [3].

Our previous study demonstrated that isoflurane reduced CD4□ Th cell frequency during the intraoperative phase compared to propofol in perioperative breast cancer patients [4]. Therefore, we aimed to elucidate the mechanistic circuitry for the isoflurane-associated reduction in CD4□ Th cell frequency. A decline in CD4□ Th cell frequency may result from decreased proliferation or increased cell death, or both, during the intraoperative period [5].

Apoptosis proceeds via extrinsic or intrinsic pathways [6]. Sevoflurane and isoflurane induced apoptosis in Jurkat T cells and human CD3 T lymphocytes through mitochondrial disruption and activation of caspase-3, independent of death receptor signaling [7]. Though propofol reduced lymphocyte expression of both FS-7 cell-associated surface antigen (Fas)/ Fas ligand (FasL) and B-cell lymphoma-2 (Bcl-2), it did not trigger apoptosis [8]. It is well established that postoperative immunosuppression following major surgery and anaesthesia manifests as peripheral T cell lymphopenia alongside leucocytosis [9,10]. Despite this, relatively few studies have explored anaesthesia-induced CD4□ Th cell death.

Building on this rationale, the present study investigated the comparative effect of isoflurane and propofol on the apoptotic regulation of perioperative breast cancer patient-derived CD4□ Th cells and Jurkat T cells.

## Materials and Methods

### Study Setting

The study included patients receiving isoflurane and propofol (n=15 per group) from the previous clinical trial cohort [Clinical Trials Registry-India (CTRI/2020/11/028826 dated Nov 3, 2020)], approved by the Institutional Ethics Committee [4].

### Magnetic Sorting of CD4^+^ Th Cells

Peripheral blood mononuclear cells (PBMCs) were extracted from whole blood at three timepoints {preoperative (1 day before surgery), intraoperative (1 h after surgical incision) and postoperative (48 h after surgery)} utilizing HiSep™ LSM 1077 (HiMedia Laboratories Pvt. Ltd., Mumbai, India). CD4^+^ Th cells were negatively isolated from PBMCs using the MojoSort™ Human T Cell Isolation Kit (BioLegend, San Diego, CA, USA) and magnetic activated cell sorting columns (Miltenyi Biotec, North Rhine-Westphalia, Germany). The purity of the sorted CD4^+^ Th cells was checked by flow cytometry (Fig. S1).

### Cell Culture and Treatment

Jurkat E6.1 cells were cultured in RPMI-1640 with 10% fetal bovine serum (FBS) and 1% penicillin-streptomycin, under a humidified environment with 5% CO_2_ at 37°C.

Due to the differences in the physicochemical properties and pharmacodynamics of volatile and intravenous anaesthetics, it was not possible to establish direct molar equivalence. Therefore, we chose concentration ranges that reflected clinically relevant exposure for each anaesthetic agent. Jurkat T cells (2 × 10□ cells/mL) were subjected to already established clinically relevant concentrations of isoflurane (Cayman, Ann Arbor, MI, USA)-0.3-1 mM (0.5-1.5 minimum alveolar concentration in humans) [11–17] and propofol (Merck-Sigma-Aldrich, Burlington, MA, USA) - 10 to 50 µM [11,18–20] for 1, 3, and 6 h. Isoflurane and propofol were dissolved in dimethyl sulfoxide (DMSO) according to the manufacturer’s protocol. Isoflurane was administered rapidly to culture plates and was promptly sealed to reduce evaporative loss and preserve liquid-phase exposure conditions. Vehicle control (0.1% DMSO) was checked to ensure the accuracy of the effect (Fig. S2). All the *in vitro* experiments were conducted in triplicate.

### Cell Viability Assay

Jurkat T cells were treated with isoflurane (0.3-1 mM) and propofol (10 to 50 µM) for 1, 3, and 6 h. Further, cells were incubated with 3-4,5-dimethylthiazol-2-yl)-2,5-diphenyl-2H-tetrazolium bromide (MTT) solution (0.5 mg.mL^-1^) for 4 h, and absorbance was detected at 570 nm using a microplate reader (Infinite M200, Tecan, Männedorf, Switzerland).

### Detection of Apoptosis

Cells were incubated with fluorescein isothiocyanate (FITC)-conjugated Annexin V and propidium iodide (PI, 5 µL) in the presence of Annexin V binding buffer for 20 min. Subsequently, binding buffer (200 µL) was added and samples were acquired on a flow cytometer [BD LSRFortessa cell analyzer; software BD FacsDiva 9.0.1, Franklin Lakes, NJ, USA] using FITC and PE channels. Data were analyzed using FlowJo version 10.8.1 [BD, Franklin Lakes, NJ, USA]. To assess caspase-dependent apoptosis, cells were pre-incubated with the caspase-3/7 inhibitor [Z-Asp-Glu-Val-Asp(OMe)-fluoromethylketon (Z-DEVD-FMK), 20 µM], [21] for 1 h before treatment with anaesthetics.

### Measurement of Mitochondrial Membrane Potential

Mitochondrial membrane potential (ΔΨm) of cells was assessed using 5,5,6,6’-tetrachloro-1,1’,3,3’ tetraethylbenzimi-dazoylcarbocyanine iodide (JC-1, Elabscience, Houston, TX, USA), acquired on a flow cytometer, as mentioned before, in the FITC and PE channels. Gating and compensation were done according to the manufacturer’s protocol.

### Cytochrome c Release Assay from Mitochondria to Cytosol

Cytochrome c release assay from mitochondria to cytosol required isolation of mitochondrial and cytosolic fractions and was followed according to Dimauro et al 2012 [22]. For the assay, cytochrome c expression was checked along with mitochondrial and cytosolic housekeeping proteins, cytochrome c oxidase (COX) IV and β-actin, respectively, using Western blotting.

### Detection of Caspase-3/7 Activity

Sorted CD4□ Th cells and Jurkat T cells were incubated with the CellEvent™ caspase-3/7 green detection reagent (Invitrogen, Waltham, MA, USA, 5 µM) [23] for 30 min in the dark at 37 °C according to the manufacturer’s instructions. Subsequently, cells were analyzed by a flow cytometer in the FITC channel. Caspase-3/7 activity was evaluated both in the presence/absence of their specific inhibitor, Z-DEVD-FMK.

### Analysis of DNA Fragmentation

DNA fragmentation was evaluated using the terminal deoxynucleotidyl transferase dUTP nick end labeling (TUNEL) assay kit according to the manufacturer’s protocol (Elabscience, Houston, TX, USA), analyzed using a flow cytometer in the FITC channel.

### Analysis of DNA Damage by Comet Assay

DNA strand breaks were checked by comet assay [24] according to our previous publication [25]. Images were obtained using a confocal microscope (FLUOVIEW FV4000, Evident-Olympus, Tokyo, Japan). The comet tail length, tail moment, and olive tail moment were measured utilizing the plugin OpenComet in Fiji ImageJ software [NIH, Bethesda, MD, USA].

### Detection of Reactive Oxygen Species (ROS) Generation

Cells were incubated with 2′-7′-dichlorodihydrofluorescein diacetate (DCFH-DA, 2 µM) at room temperature for 10 min and analyzed by flow cytometry in the FITC channel. ROS production was also assessed in the presence/absence of the negative control N-acetyl cysteine (NAC). Jurkat T cells were pretreated for 1 h with NAC (5 mM) [26] before treatment with anaesthetics.

### Detection of T Cell Proliferation

Jurkat T cells were fixed in 2% paraformaldehyde for 15 min, permeabilized with 0.2% Triton X-100, and blocked with 1% bovine serum albumin (BSA) for 1 h. Cells were incubated overnight at 4°C with Ki-67□ primary antibody, washed, and stained with FITC-conjugated secondary antibody for 2 h. Ki-67 cells were analyzed by flow cytometer.

Jurkat T cells were labeled with carboxyfluorescein succinimidyl ester (CFSE), treated with anaesthetics for 3 h, and further cultured in drug-free medium for 24-48 h. Data were acquired in the FITC channel in a flow cytometer. Gating was done based on the fixed samples (stained sample was fixed in 2 % paraformaldehyde at the beginning of the experiment).

### Reverse transcription real-time quantitative polymerase chain reaction (RT-qPCR)

The gene expression was analyzed by RT-qPCR according to our previous publication [25]. Gene expression was analyzed using Genious 2× SYBR Green Fast qPCR Mix and gene-specific primers (Table S1) on a LightCycler® 96 (Roche Diagnostics, Rotkreuz, Switzerland). Gene expression analysis included comparison between intraoperative or postoperative and preoperative transcript profile in sorted CD4^+^ Th cells, and treated versus control in Jurkat T cells. Relative fold changes were calculated by 2^−ΔΔCt^ method with normalization to glyceraldehyde-3-phosphate dehydrogenase.

### Western Blot

Protein samples (50 µg each) from Jurkat T cells were resolved on sodium dodecyl sulfate-polyacrylamide gel electrophoresis and transferred onto polyvinylidene fluoride membranes. Membranes were blocked with 5% BSA/ Tris-buffered saline with 0.1% Tween® 20, incubated overnight with primary antibodies [anti-human Bcl-2 antagonist/killer (Bak), Bcl-2, myeloid cell leukemia-1 (Mcl-1), caspase-3, active caspase-3, poly(ADP-ribose) polymerase 1 (PARP1), FasR, caspase-8, c-Jun N-terminal kinases 1/2/3 (JNK)1/2/3, phospho (p)-JNK1/2/3-T183/T183/T221, extracellular signal-regulated kinase (ERK)1/2, p-ERK1-T202/Y204 + ERK2-T185/Y187, p38 mitogen-activated protein kinase (MAPK), p-p38 MAPK-T180/Y182, and β-actin] at 4°C. Further, membranes were incubated with horseradish peroxidase (HRP)-conjugated secondary antibodies for 2 h. Bands were developed using enhanced chemiluminescence super kit (Abclonal Woburn, MA, USA), and observed under gel documentation system (iBright FL1500 Imaging System, Thermo Fisher Scientific, Waltham, MA, USA). Band intensities were measured using ImageJ software (NIH, Bethesda, MD, USA). For the inhibitor study, Jurkat T cells were pretreated with a selective p38 MAPK inhibitor (SB203580; 10 µM) [27] for 1h before treatment with anesthetics.

### Enzyme-linked Immunosorbent Assay (ELISA)

Protein samples (10 µg) extracted from sorted CD4^+^ Th cells were used for ELISA according to our previous publication [25]. Cells were incubated with primary antibodies anti-human Bcl-2, Mcl-1, Bcl-2 associated x (Bax), active caspase-3, caspase-3, PARP1, and β-actin (1:500 dilution) for 2 h at room temperature, followed by incubation with HRP-conjugated secondary antibody (1:1000 dilution) for 1 h. Wells were then incubated with tetramethylbenzidine substrate for 30 to 45 min until color developed, and the reaction was halted with H_2_SO_4_ (1M). Absorbance was measured at 450 nm using a microplate reader.

### Bioinformatics Analysis

Protein-chemical interactions for propofol and isoflurane were identified using the Search Tool for Interactions of Chemicals (STITCH) database (http://stitch.embl.de/) with the maximum number of interactions in the first shell set to 20 and the minimum required interaction score set to 0.700 (high confidence). Common differentially expressed genes (DEGs) regulated by both anaesthetics were retrieved from the Comparative Toxicogenomics Database (CTD, https://ctdbase.org/). Functional enrichment analysis of common DEGs was performed using ShinyGO v0.82 (https://bioinformatics.sdstate.edu/go/) for Gene Ontology (GO; cellular component, molecular function and biological process) and Kyoto Encyclopedia of Genes and Genomes (KEGG) pathway analysis.

### Statistical Analyses

A linear mixed-effects model was used to evaluate the effects of anaesthetic technique and perioperative timepoints on biomarker levels, while adjusting for clinically relevant covariates, including age, body mass index, ASA status, duration of anaesthesia, type of surgery, and disease stage. Patient ID was included as a random effect to account for within-subject correlations arising from repeated measurements, whereas anaesthetic group and timepoints were treated as fixed effects. An interaction between anaesthetic group and timepoints were incorporated to assess whether the trajectory of biomarker changes across perioperative timepoints differed between the two anaesthetic groups [28]. Graphs were plotted based on the estimated marginal means of pairwise comparisons. Propofol was considered the reference category for the anaesthetic group variable, and estimates for the isoflurane group were interpreted relative to propofol. Variance inflation factor was used to assess multicollinearity among the independent variables, and all values were below the commonly accepted threshold of 5, indicating no evidence of multicollinearity among the predictors. For all *in vitro* experiments with Jurkat T cells involving three independent groups (control, isoflurane, propofol), one-way ANOVA was performed with post hoc Bonferroni test. Data were analyzed using SPSS (IBM, ver. 25.0, Chicago, IL, USA) and R (version 4.3.3). A two-tailed p-value of <0.05 was considered statistically significant.

## Results

### Isoflurane & Propofol Differentially Modulate Apoptosis

Bioinformatics analyses were used to discern shared molecular pathways that may be influenced by both anaesthetics before conducting targeted mechanistic experiments. STITCH analysis identified that four proteins [heme oxygenase 1, protein kinase C gamma, caspase-3 (*CASP3)*, and solute carrier family 1 member 1] were targeted by both anaesthetics (Fig. S3a, Table S2). CTD analysis showed that *CASP3* activation had a stronger association with isoflurane than propofol (Fig. S3b). CTD analysis further revealed 243 DEGs linked to isoflurane and 467 to propofol, with 36 genes overlapping between the two datasets (Fig. S3c). These 36 shared genes were subjected to functional enrichment analysis using *ShinyGO v0.82* (FDR cutoff = 0.05; pathway size range = 2–5000). Top 10 enriched GO terms were ranked by fold enrichment for cellular component (Fig. S3d), molecular function (Fig. S3e), biological process (Fig. S3f) and KEGG pathways (Fig. S3g). Interestingly, apoptosis ranked as the third most enriched KEGG pathway (Fig. S3g). Collectively, these analyses highlighted apoptosis as a key cellular mechanism impacted by the anaesthetic agents.

To further substantiate these findings, we analyzed sorted CD4□ Th cells directly from pre-, intra-, and postoperative patient samples. No significant difference in clinicopathological characteristics was found between the two anaesthetic groups (Table S3). In intragroup analysis, apoptosis was significantly increased during the intra-than pre- and postoperative period of the isoflurane group (Fig. 1a, b). Intergroup analysis showed that isoflurane significantly increased apoptosis of CD4□ Th cells compared with propofol during the intraoperative period (β: 15.93; 95% CI: 5.52 to 26.35; Table S4, Fig. 1c). TUNEL^+^ cells were found to be significantly high in the intra-than pre- and postoperative period of isoflurane group and no changes were observed in propofol group (Fig. 1d, e). Intergroup analysis exhibited that TUNEL^+^ cells were significantly high with isoflurane than propofol during the intraoperative period (β: 160.47; 95% CI: 102.51 to 218.42; Fig. 1f; Table S4). Within-group comparisons demonstrated a decline in ΔΨm of CD4□ Th cells during the intra-versus pre- and postoperative period in the isoflurane group, whereas propofol was associated with no significant changes (Fig. 1g, h). Consistently, intergroup comparison revealed that ΔΨm was markedly reduced with isoflurane relative to propofol during the intraoperative period (β: -1.74; 95% CI: -2.40 to -1.07; Fig. 1i; Table S4). Isoflurane significantly induced ROS in CD4□ Th cells during the intraoperative period compared to the pre- and postoperative period, while no effect was found with propofol (Fig. 1j, k). Intergroup analysis showed increased ROS generation by isoflurane than propofol during the intraoperative period (β: 409.93; 95% CI: 225.55 to 594.31; Fig. 1l; Table S4).

**Fig. 1:**
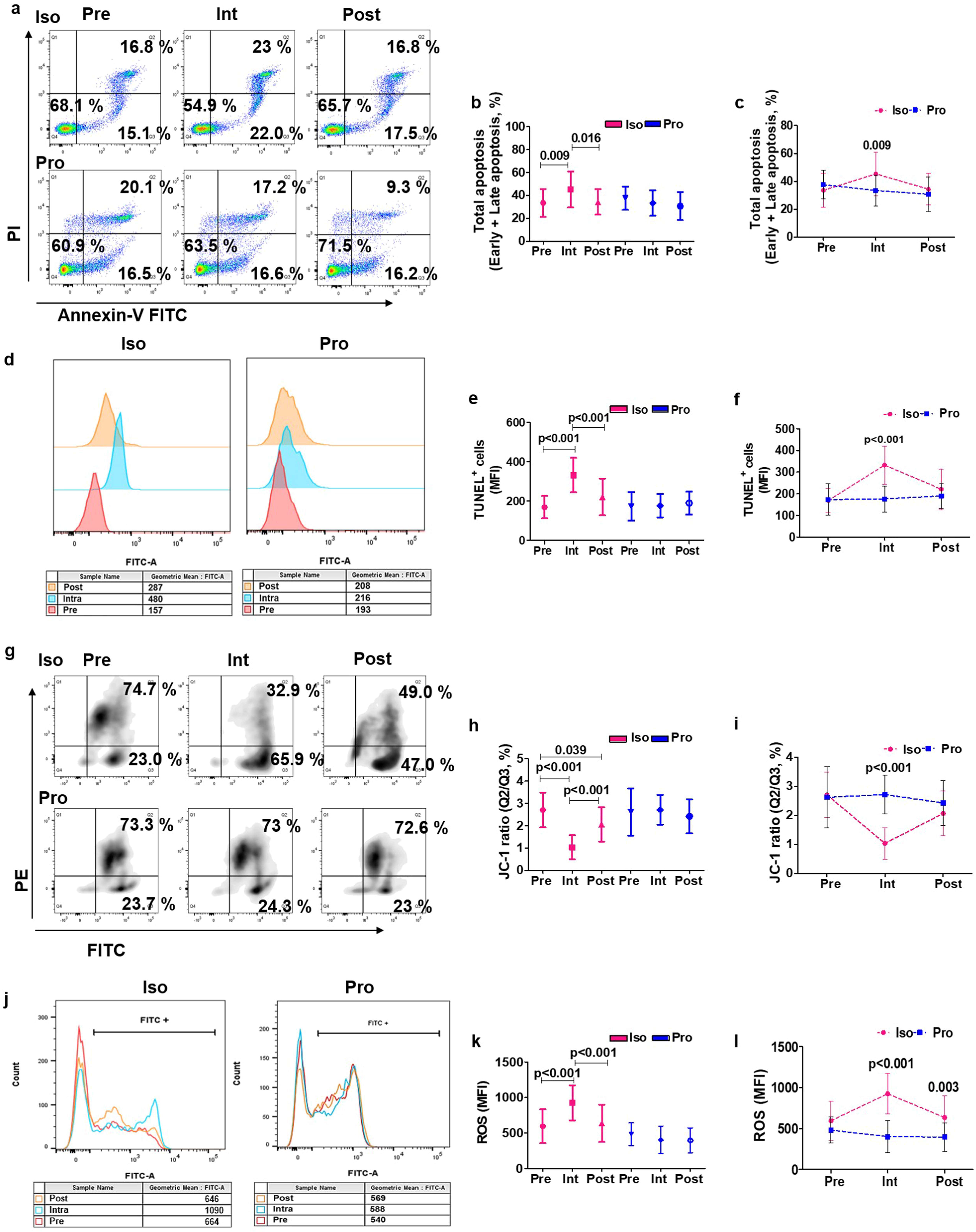
Effect of isoflurane and propofol on the apoptosis of sorted CD4□ Th cells in perioperative breast cancer patients. Representative flow cytometric analysis of sorted CD4□ Th cells in females anesthetized with isoflurane and propofol (a); Comparative influence of isoflurane and propofol on intragroup (b) and intergroup (c) analysis of total apoptotic cells; Representative flow cytometric analysis of TUNEL assay in females anesthetized with isoflurane and propofol (d); Comparative influence of isoflurane and propofol on intragroup (e) and intergroup (f) analysis of TUNEL^+^ cells; Representative flow cytometric analysis of mitochondrial membrane potential by JC-1 staining (g) in females anesthetized with isoflurane and propofol; Comparative influence of isoflurane and propofol on intragroup (h) and intergroup (i) analysis of mitochondrial membrane potential; Representative flow cytometric histogram overlay of ROS generation (j) in females anesthetized with isoflurane and propofol; Comparative effect of isoflurane and propofol on ROS production with intragroup analysis (k) and intergroup analysis (l) at different timepoints in perioperative breast cancer patients. [The graphs were plotted based on the mean ±SD; Int, Intraoperative; Iso, Isoflurane; JC-1, 5,5,6,6′-Tetrachloro-1,1′,3,3′-tetraethylbenzimidazolylcarbocyanine iodide; Post, Postoperative; Pre, Preoperative; Pro, Propofol; ROS, Reactive oxygen species; TUNEL, Terminal deoxynucleotidyl transferase dUTP nick-end labelling]

### Isoflurane & Propofol Differentially Regulated Apoptosis Signaling in CD4□ Th Cells

Apoptosis signaling in patient-derived CD4□ Th cells demonstrated that anti-apoptotic Bcl-2 and Mcl-1 showed no differences between the anaesthetic groups (Fig. 2a, b). While, pro-apoptotic Bax and activated caspase-3 were elevated during the intra-compared with pre- and postoperative phases of the isoflurane group (Fig. 2c, d). Intergroup analysis revealed overexpression of Bax (β: 0.38; 95% CI: 0.23 to 0.54; Table S4) and activated caspase-3 (β: 0.35; 95% CI: 0.15 to 0.54; Table S4) by isoflurane than propofol during the intraoperative period (Fig. 2c, d). Expression of PARP1 was elevated in the intraoperative than pre- and postoperative period of isoflurane group (Fig. 2e). Isoflurane increased caspase-3/7 activity during the intra- and postoperative periods compared to preoperative baseline, whereas propofol showed reduced caspase-3/7 activity in the post-compared to pre- and intraoperative period (Fig. 2f, g). Intergroup comparisons confirmed higher caspase-3/7 activity with isoflurane than propofol during the intra-(β: 1335.93; 95% CI: 1046.79 to 1625.07; Table S4) and postoperative period (β: 1514.93; 95% CI: 1225.79 to 1804.07; Fig. 2g; Table S4). *BCL2*, *MCL1* and *PARP1* transcripts remained unaffected by anaesthetics. Whereas, *BAX* and *CASP3* were significantly elevated by isoflurane than propofol during the intraoperative period (Fig. 2h).

**Fig. 2:**
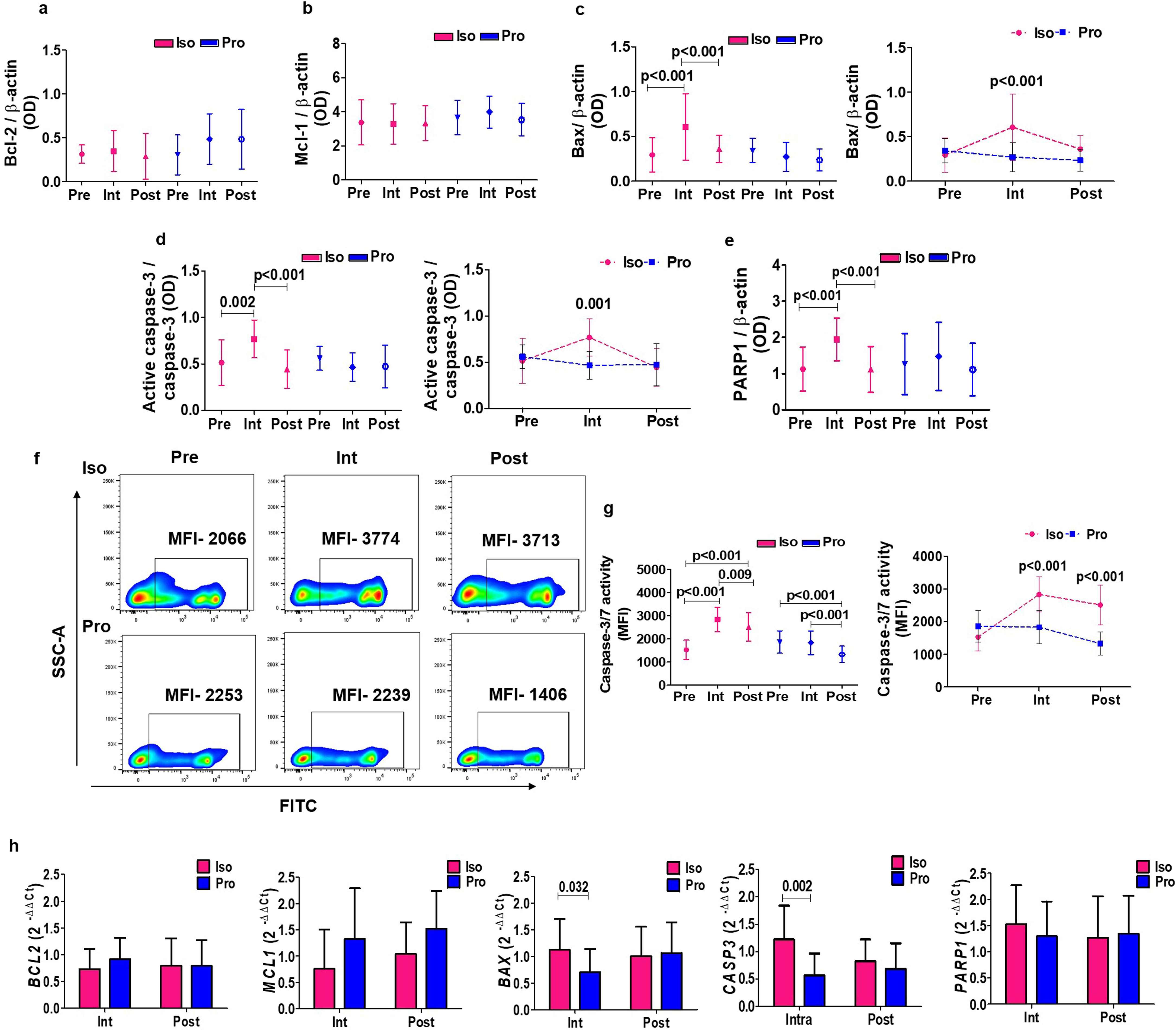
Effect of isoflurane and propofol on apoptosis signaling in sorted CD4□ Th cells of perioperative breast cancer patients. Comparative effect of isoflurane and propofol on the protein expression levels with intragroup analysis of Bcl-2 (a), intragroup of Mcl-1 (b), intra- and intergroup of Bax (c), intra- and intergroup of active Caspase-3 (d) and intragroup of PARP1 (e); Representative flow cytometric analysis of Caspase-3/7 activity (FITC+) in sorted CD4^+^ Th cells (f) of females anesthetized with isoflurane and propofol; Comparative influence of isoflurane and propofol on intra- and intergroup (g) analysis of Caspase-3/7 activity; Comparative influence of isoflurane and propofol on the gene expression levels of *BCL2, MCL1, BAX, CASP3* and *PARP1* expression during intra-and postoperative period compared to preoperative period in the breast cancer patients (h) at different timepoints in perioperative breast cancer patients. [The graphs were plotted based on the mean ±SD; Bax, Bcl-2 antagonist/killer; Bcl-2, B-cell lymphoma-2; Int, Intraoperative; Iso, Isoflurane; Mcl-1, Myeloid cell leukemia-1; PARP1, Poly (ADP-ribose) polymerase 1; Post, Postoperative; Pre, Preoperative; Pro, Propofol]

### Effect of Isoflurane and Propofol on Mitochondria-Dependent Apoptosis

To validate the clinical findings independent of surgical influence and other perioperative drugs, Jurkat T cells were treated with isoflurane (0.3-1 mM) [11–17] and propofol (10-50 μM) [11,18–20]. Isoflurane (1 mM) for 3 h and 6 h caused a reduction in cell viability, whereas propofol had no comparable effect (Fig. S4a). As patient exposure was limited to 3 h, subsequent experiments were standardized to this duration. Apoptosis was evaluated using isoflurane (0.3-1 mM) and propofol (10-20 μM) for 3 h (Fig. S4b). Since cell viability was similar between 20 μM and 50 μM propofol, the higher concentration was excluded. Total apoptosis (early + late) was significantly increased with 1 mM isoflurane compared to both control and 20 μM propofol (Fig. S4c). Accordingly, all subsequent experiments were performed using isoflurane (1 mM) and propofol (20 μM) for 3 h.

Isoflurane-induced apoptosis was not mediated by extrinsic pathway as FasR and caspase-8 protein/RNA profile exhibited insignificant changes between anaesthetic groups (Fig.3a, b). Isoflurane significantly reduced the JC-1 ratio compared with control and propofol, indicating ΔΨm loss and supporting mitochondria-mediated apoptosis (Fig. 3c). Intrinsic pathway analysis showed isoflurane significantly increased Bak, reduced Bcl-2 expression relative to both control and propofol but Mcl-1 showed no change between either anaesthetic (Fig. 3d). Isoflurane also reduced pro caspase-3 and increased active caspase-3 and PARP1 expression compared to control and propofol (Fig. 3d). *BAK1*, *CASP3* and *PARP1* were upregulated while *BCL2* was downregulated by isoflurane than propofol (Fig. 3e). *MCL1* did not show any alteration between the groups (Fig. 3e). Consistently, isoflurane increased cytochrome c release from mitochondria to cytosol (Fig. 3f, g) which might have triggered intrinsic pathway of apoptosis by activating caspase-3/7 activity (Fig. 3h). Further, isoflurane induced caspase-3/7 activity (Fig. 3i) and corresponding apoptosis (Fig. 3j) was significantly reduced in comparison to propofol with caspase-3/7 specific inhibitor Z-DEVD-FMK.

**Fig. 3:**
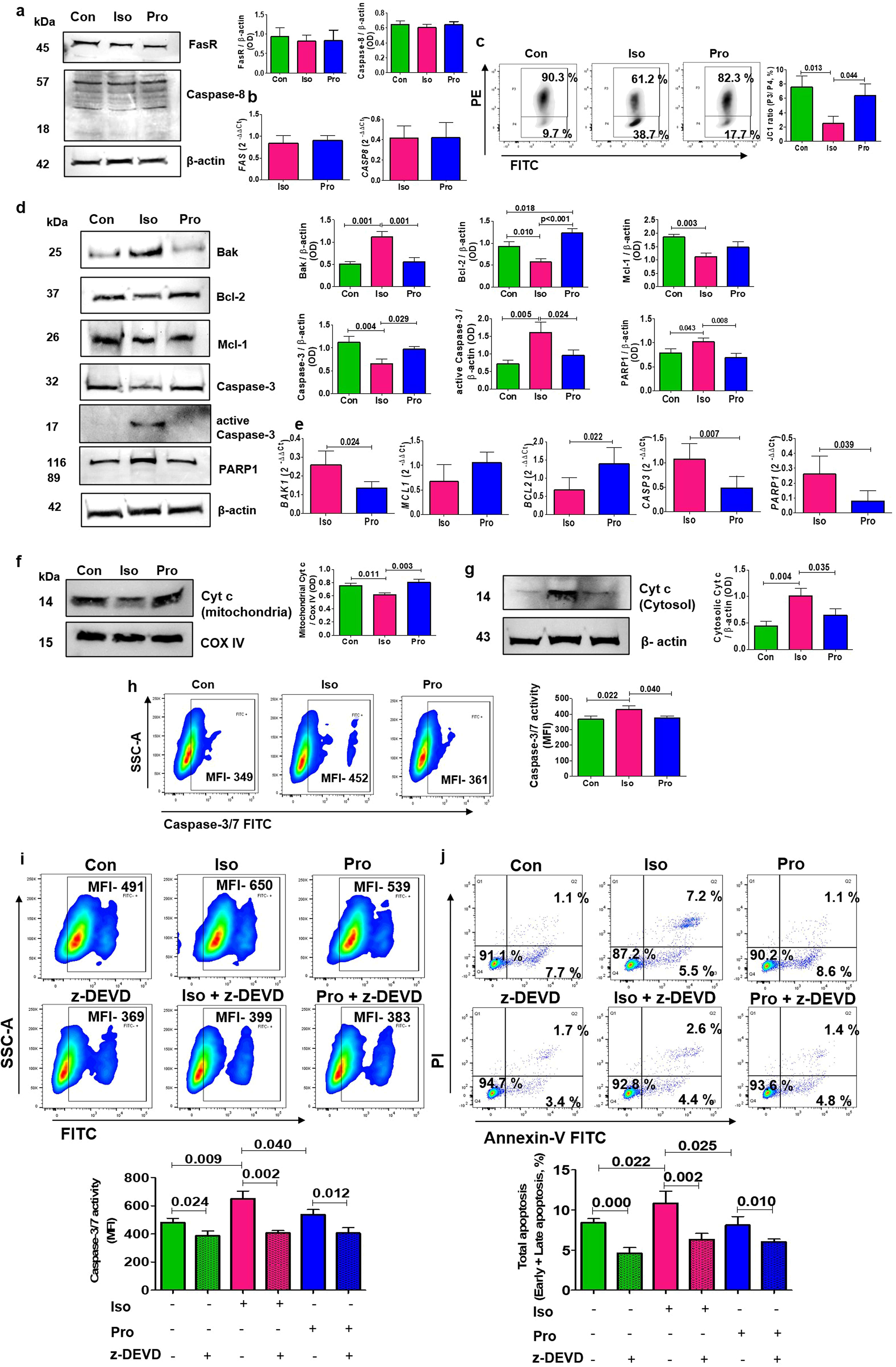
Effect of isoflurane and propofol on apoptosis signaling in Jurkat T cells. Comparative effect of isoflurane and propofol on extrinsic apoptotic pathway as observed with protein expression of FasR and Caspase-8 (a) and RNA expression of *FAS* and *CASP8* (b); Comparative effect of isoflurane and propofol on mitochondrial membrane potential of Jurkat T cells as evidenced from representative flow cytometry with JC-1 and JC-1 ratio (P3/P4, c); Comparative effects of isoflurane and propofol on the intrinsic pathway apoptotic molecules as evident from representative western blots and mean band intensities of Bak, Bcl-2, Mcl-1, Caspase-3, active Caspase-3, PARP1 and β-actin (d); Gene expression analysis of *BAK, MCL1, BCL2, CASP3* and *PARP1* in terms of fold change with respect to the control samples (e); Representative western blots of Cyt c expressed in mitochondrial (f) and cytosolic fractions (g) and their comparative band intensities in isoflurane and propofol treated Jurkat T cells; Representative flow cytometric analysis of Caspase-3/7 activity and the comparative effect of isoflurane and propofol on the same (h); Effect of isoflurane and propofol on Caspase-3/7 activity in presence and absence of Z-DEVD-FMK as evident from representative flow cytometric analysis and comparative plots (i); Effect of isoflurane and propofol on apoptosis in presence and absence of Caspase-3/7 specific inhibitor, Z-DEVD-FMK, as evident from representative flow cytometric quadrant plots with annexin V and PI and comparative plots (j). [The graphs were plotted based on the mean± SD; Bax, Bcl-2-associated X protein; Bcl-2, B-cell lymphoma-2; Con, Control; Cyt c, Cytochrome c; COX IV, Cytochrome c oxidase; FAS, Fas cell surface death receptor; Iso, Isoflurane; JC-1, 5,5,6,6′-tetrachloro-1,1′,3,3′-tetraethylbenzimidazolylcarbocyanine iodide; Mcl-1, Myeloid cell leukemia-1; MFI, Mean fluorescence intensity; PARP1, Poly (ADP-ribose) polymerase 1; Pro, Propofol; Z-DEVD-FMK, Z-Asp-Glu-Val-Asp(OMe)-fluoromethylketone]

### Isoflurane Triggered DNA Damage and ROS Generation

Isoflurane increased TUNEL^+^ cells (apoptotic DNA fragmentation) compared with control and propofol in Jurkat T cells (Fig. 4a). Isoflurane, but not propofol, increased DNA damage as evident from comet tail length, tail moment, and olive tail moment (Fig. 4b).

**Fig. 4:**
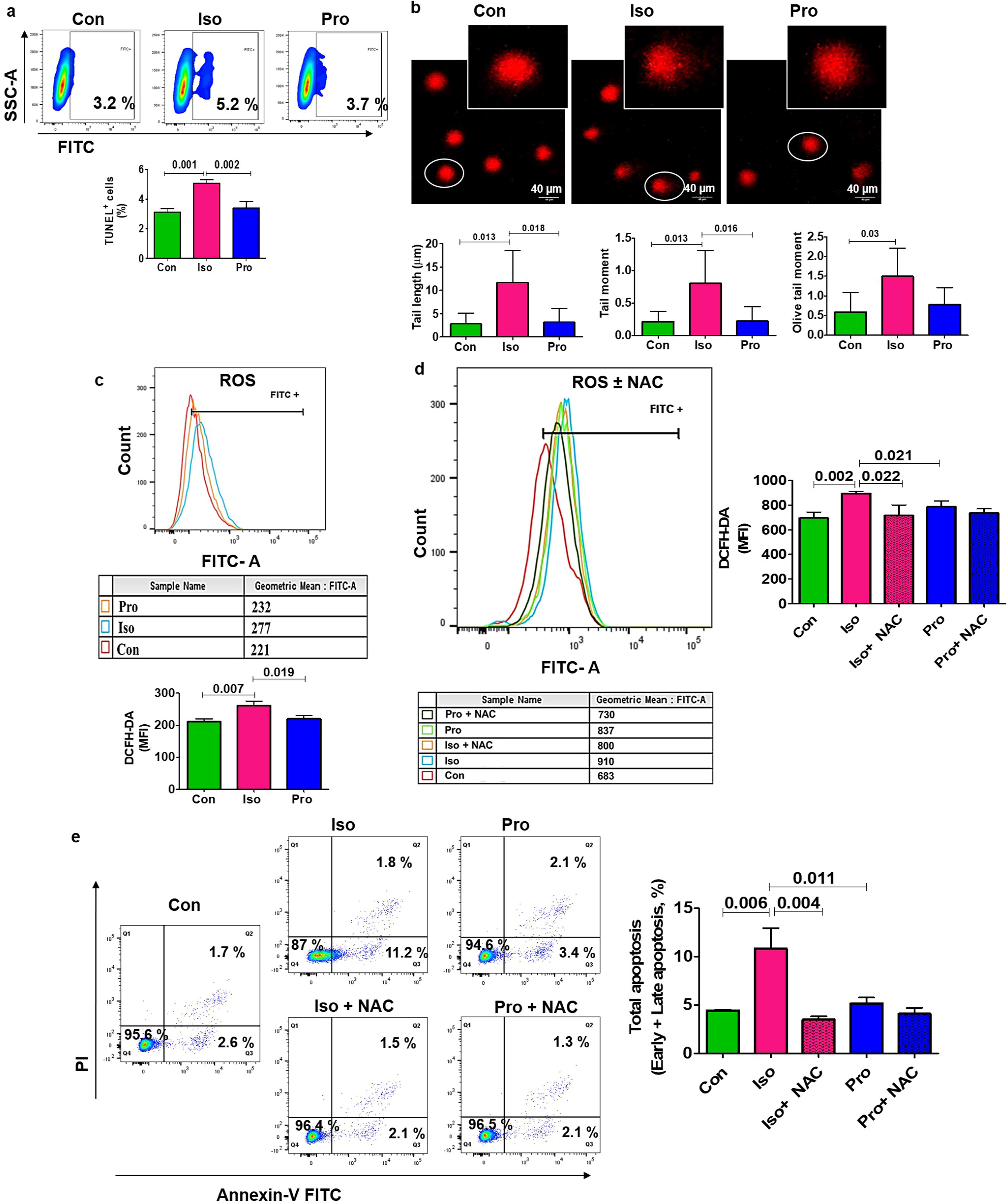
Effect of isoflurane and propofol on DNA damage and ROS generation in Jurkat T cells. Flow cytometric representation of TUNEL assay and comparative effect of isoflurane and propofol on TUNEL(+) cells (a); Effect of isoflurane and propofol on DNA damage by comet assay in Jurkat T cells under 40× magnification and comparative analysis of comet tail length, tail moment and olive moment (b); Effect of isoflurane and propofol on ROS generation using DCFH-DA as depicted by representative flow cytometric histogram overlay and comparative MFI (c); Flow cytometric histogram overlay of ROS generation with isoflurane and propofol in presence and absence of NAC and their comparative analysis (d); Flow cytometric apoptosis analysis with isoflurane and propofol in presence and absence of NAC and their comparative effects on the total apoptosis (e). [The graphs were plotted based on the mean± SD; Con, Control; DCFH-DA, 2’,7’-dichlorodihydrofluorescein diacetate; Iso, Isoflurane; MFI, Mean fluorescence intensity; NAC, N-acetylcysteine; Pro, Propofol; ROS, Reactive oxygen species; TUNEL, Terminal deoxynucleotidyl transferase dUTP nick-end labeling].

Since ROS are known mediators of apoptosis, we evaluated intracellular ROS levels, which increased significantly with isoflurane compared to control and propofol (Fig. 4c). The antioxidant NAC effectively suppressed isoflurane-induced ROS generation (Fig. 4d) and apoptosis (Fig. 4e) in Jurkat T cells, supporting that isoflurane-induced apoptosis was ROS dependent.

### Impact on Apoptosis and Cell Proliferation Under Drug-Free Condition

To evaluate post-treatment apoptotic effects, Jurkat T cells were cultured in drug-free medium following anaesthetic exposure (3 h). After 24 h (Fig. 5a, b) and 48 h (Fig. 5c, d) in drug-free medium, apoptosis returned towards baseline, indicating that isoflurane removal restored T cell viability. This observation paralleled the postoperative recovery of apoptosis in CD4^+^ Th cells from perioperative breast cancer patients.

**Fig. 5:**
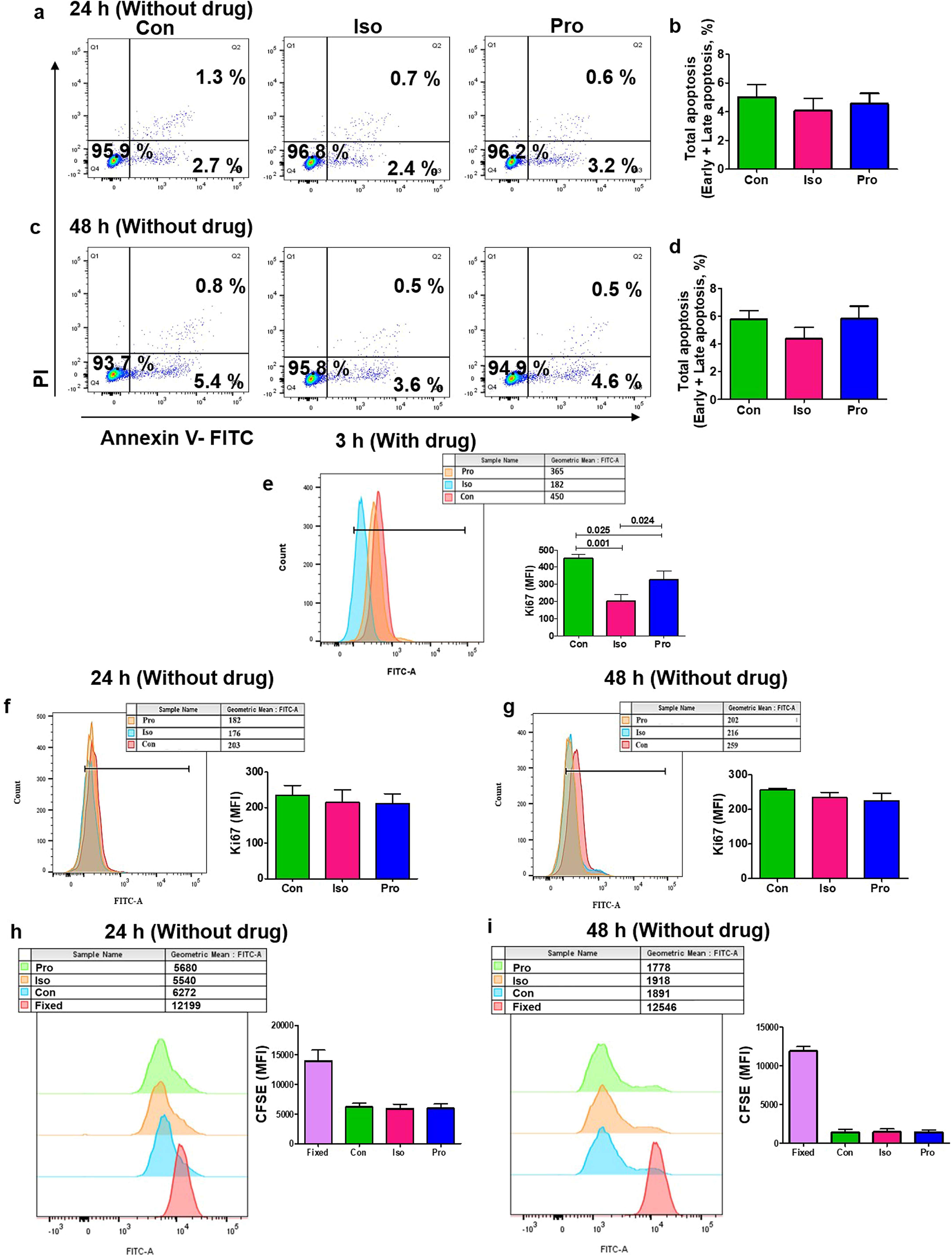
Effect on apoptosis and proliferation of Jurkat T cells in drug free condition. Representative flow cytometry and comparative effect of isoflurane and propofol on total apoptosis after a 3 h initial treatment, followed by analysis in drug-free media at 24 h (a, b) and 48 h (c, d) in Jurkat T cells; Representative flow cytometric histogram overlay and comparative effect of isoflurane and propofol on the Ki67 expression after 3 h treatment (e); Representative flow cytometric histogram overlays and corresponding graphs of MFI showing the comparative effects of isoflurane and propofol on Ki67 expression in Jurkat T cells following 3 h of initial treatment, with subsequent monitoring in drug-free media up to 24 h (f) and 48 h (g); Representative flow cytometric histogram overlays and corresponding graphs depicting the differential effects of isoflurane and propofol on CFSE in Jurkat T cells after a 3 h initial treatment, followed by analysis in drug-free media at 24 h (h) and 48 h (i); [The graphs were plotted based on the mean± SD; CFSE, Carboxyfluorescein succinimidyl ester; Con, Control; Iso, Isoflurane; MFI, Mean Fluorescence Intensity; Pro, Propofol]

To assess anaesthetic effects on the proliferation of Jurkat T cells, Ki67 expression was measured after treatment with isoflurane/propofol (3h). Ki67 expression was significantly reduced by both anaesthetics compared with controls (Fig. 5e). However, propofol-treated cells showed relatively higher Ki67 expression than isoflurane-treated cells (Fig. 5e). Under drug-free conditions, no significant differences were observed for Ki67 expression between groups after 24 h (Fig. 5f) or 48 h (Fig. 5g). Similarly, CFSE staining revealed no difference in proliferation between groups after 24 h (Fig. 5h) or 48 h (Fig. 5i).

### Isoflurane & Propofol Altered MAPK Signaling in Jurkat T cells

To further elucidate the upstream signaling underlying anaesthetic agent-induced apoptosis in Jurkat T cells, we examined the MAPK signaling pathways (JNK, p38, and ERK). The relative expression of p-ERK1-T202/Y204 + ERK2-T185/Y187 to total ERK1/2 was significantly reduced with isoflurane than control and propofol (Fig. 6a, b). However, the relative expression of p-p38 MAPK-T180/Y182 to total p38 MAPK significantly increased with isoflurane compared to control (Fig. 6a, b). However, relative expression of p-JNK1/2/3-T183/T183/T221 to total JNK1/2/3 showed a similar significant reduction with isoflurane and propofol compared to control (Fig. 6a, b). *MAPK1* (encoded for ERK2), *MAPK9* (encoded for JNK2) and *MAPK14* (encoded for p38α) did not show change at the transcript level by these anaesthetics (Fig. 6c). Isoflurane induced p38 MAPK phosphorylation compared to control and propofol was further reduced with p38 MAPK inhibitor SB203580 (Fig. 6d, e). Increased p38 MAPK activity might be linked with increased apoptosis in a ROS dependent manner. We observed that isoflurane-associated apoptosis reduced significantly with p38 MAPK inhibitor (Fig. 6f, g).

**Fig. 6:**
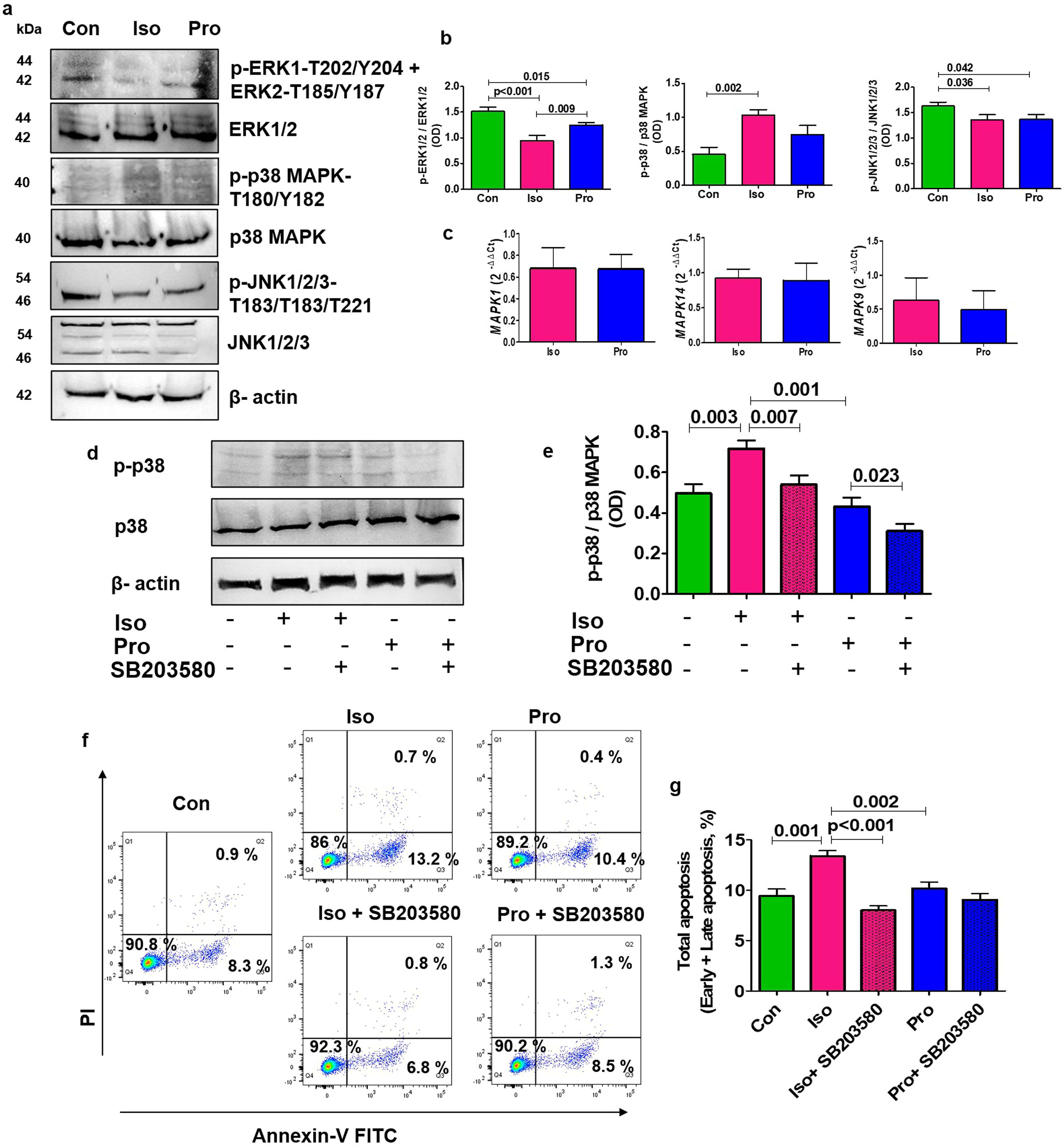
Effect of isoflurane and propofol on the MAPK signaling pathway in Jurkat T cells. Effect of isoflurane and propofol on MAPK pathway molecules-p-ERK1-T202/Y204 + ERK2-T185/Y187, ERK1/2, p-p38 MAPK-T180/Y182, p38 MAPK, p-JNK1/2/3-T183/T183/T221, JNK1/2/3 and β-actin as evidenced by representative western blots (a); and comparative band intensities (b); Effect of isoflurane and propofol on gene expression of *MAPK1*, *MAPK14* and *MAPK9* (c) in Jurkat T cells; Representative Western blots of p-p38 MAPK-T180/Y182 depicting effect of isoflurane and propofol with/without p-p38 inhibitor-SB203580 (d); Comparative influence of isoflurane and propofol in presence/absence of SB203580 on relative expression of p-p38 to total p38 MAPK (e); Representative flow cytometry (f) and comparative influence of isoflurane and propofol in presence/absence of SB203580 on the apoptosis of Jurkat T cells (g). [The graphs were plotted based on the mean± SD; Con, Control; ERK, Extracellular-signal-regulated kinase; Iso, Isoflurane; JNK, c-Jun N-terminal kinases; MAPK-1, -14, -19, Mitogen-activated protein kinase -1, -14, -19; Pro, Propofol]

## Discussion

The selection of intravenous over volatile anaesthetics during cancer surgery has been a subject of debate regarding their immunosuppressive effects [3,11]. In our previous study, we observed that isoflurane relative to propofol, reduced the intraoperative frequency of CD4^+^ Th cells in perioperative breast cancer patients [4]. This prompted us to investigate the probable reasons for such a reduction. Prior studies reported that isoflurane activated ROS-dependent apoptosis in neuroglioma cells and mouse brain tissue [29], disrupted mitochondrial function [30] and promoted oxidative stress and DNA damage in human neuroglioma cells [31]. Repetitive exposure of mice with dexmedetomidine and propofol increased apoptosis of CD4^+^ T cells [32]. By integrating *ex vivo* and *in vitro* analyses, this study delineated the differential apoptotic effects of isoflurane and propofol on CD4^+^ Th cells.

Bioinformatic analysis propagated apoptosis as one of the important cellular pathways regulated by isoflurane and propofol. In consilience with the bioinformatic analysis and the earlier reports, we observed that isoflurane significantly increased CD4□ Th cell apoptosis during the intraoperative period, in contrast to propofol, and similar effects were observed in Jurkat T cells.

Isoflurane exhibited reduction in mitochondrial membrane potential and increased TUNEL positivity, confirming activation of mitochondria-mediated apoptotic pathways in perioperative breast cancer patient-derived CD4□ Th cells and Jurkat T cells. Isoflurane induced upregulation of pro-apoptotic Bax in sorted CD4^+^ Th cells. On the other hand, isoflurane increased Bak and PARP1 and reduced Bcl-2 in Jurkat T cells. Moreover, isoflurane induced cleavage of caspase-3 in both sorted CD4^+^ Th cells and Jurkat T cells. Isoflurane also increased the release of cytochrome c in the cytosol from mitochondria in Jurkat T cells. Evidence of isoflurane-induced apoptosis and caspase-3/7 activity in sorted CD4^+^ Th cells were mirrored in Jurkat T cells. Further use of caspase-3/7 specific inhibitor Z-DEVD-FMK confirmed that isoflurane triggered a mitochondrial, caspase-3/7-dependent intrinsic pathway in Jurkat T cells. Expression of caspase-8 and FasR remained unchanged, suggesting apoptosis was independent of the extrinsic pathway. We observed that isoflurane increased ROS production in both sorted CD4□ Th cells and Jurkat T cells compared to propofol. Pretreatment with NAC effectively attenuated ROS accumulation and prevented isoflurane-induced apoptosis in Jurkat T cells. These findings aligned with a previous report that associated isoflurane with enhanced ROS generation in neuronal cells [29]. Collectively, the results indicated that isoflurane-induced apoptosis in CD4□ Th cells and Jurkat T cells was primarily ROS-dependent and mitochondria-driven caspase-3/7 activation. This was consistent with earlier studies, which reported that isoflurane triggered mitochondrial dysfunction, cytochrome c release, and caspase-9 activation in neuronal cells [29] and induced apoptosis through oxidative stress and MAPK perturbation [33].

Further, we examined key signaling mediators of cell survival and death, mainly the MAPK (ERK, p38, JNK) signaling pathways [34]. In Jurkat T cells, isoflurane treatment led to downregulation of p-ERK1/2 and upregulation of p-p38 MAPK, while p-JNK1/2/3 levels remained unaltered. Neither anaesthetic significantly altered *MAPK1*, *MAPK9*, or *MAPK14* expression in Jurkat cells, suggesting post-transcriptional or post-translational regulation of MAPK signaling.

Additionally, isoflurane induced apoptosis through activation of p-p38 MAPK signaling, both of which were downregulated by p38 MAPK inhibitor in Jurkat T cells. These observations were in concordance with earlier reports which stated that isoflurane induced p38 MAPK expressions in neuronal or endothelial cells [35,36].

This apoptotic effect appeared to be exposure-dependent, as it reversed within 24-48 h upon isoflurane withdrawal from Jurkat T cells, aligning with reduced postoperative apoptosis and recovery of CD4^+^ Th-cell frequencies in patient samples.

Long-term clinical impact of anaesthetics has been an area of heterogeneous evidence: some studies reported better recurrence-free survival with propofol compared to volatile agents [37,38], while others observed no significant differences in progression-free survival between intravenous and volatile anaesthetics [39,40]. The isoflurane-associated transient immune disturbances during the intraoperative phase may facilitate micrometastasis and poor prognosis. Notably, in the long-term follow-up of our previous study [4], the isoflurane group recorded two recurrences and two fatalities, while propofol group exhibited no such incidents. However, due to the small cohort size, these findings may be interpreted to generate hypotheses. Future large-scale prospective studies incorporating perioperative immune biomarkers alongside long-term oncological results may ascertain whether the transient immune modulation associated with anaesthetics has significant clinical implications.

Study limitations included potential evaporative loss of isoflurane with liquid phase exposure, though conditions were standardized across experiments. Additionally, the use of Jurkat E6.1 cells did not fully reflect primary T cell biology, but enabled direct assessment of drugs without systemic influences.

Therefore, through integrated analyses of patient-derived CD4^+^ Th cells and mechanistic validations in Jurkat T cells, this study identified the ROS-p38-caspase-3/7 signaling axis and the reversible nature of isoflurane-induced apoptosis. In contrast, propofol exerted a protective, anti-apoptotic influence by maintaining mitochondrial integrity.

## Acknowledgments

The authors are grateful to Dr. Jayanta Chakrabarti, Director, Chittaranjan National Cancer Institute, for providing the project grant, fellowship to PS, and all infrastructural facilities. The authors subsequently reviewed and edited the content and take full responsibility for the final published version of the manuscript. The study included patients from a clinical trial cohort-Clinical Trials Registry-India (CTRI/2020/11/028826 dated Nov 3, 2020; PI name: Dr. Deepa Chakrabarti). The authors declare no competing interests.

