## Supplementary Materials for "Differential Impact of Isoflurane and Propofol on Apoptotic Regulation of Helper T cells"

***Enrichment of CD4⁺ Th cells from Blood Samples of Perioperative Breast Cancer Patients***

CD4^+^ Th cells were negatively isolated from the peripheral blood of breast cancer patients at three timepoints- preoperative, intraoperative and postoperative samples. Further, purity of the isolated samples was checked by flow cytometry. The flow cytometric data of the represented sample showed 50.8 % of CD3^+^CD4^+^CD8^-^ Th cells before sorting (Fig. S1a) and after sorting the purity of CD3^+^CD4^+^CD8^-^ Th cells reached to 97.6 % (Fig. S1b). Further analysis was carried out with the sorted CD4^+^ Th cells.

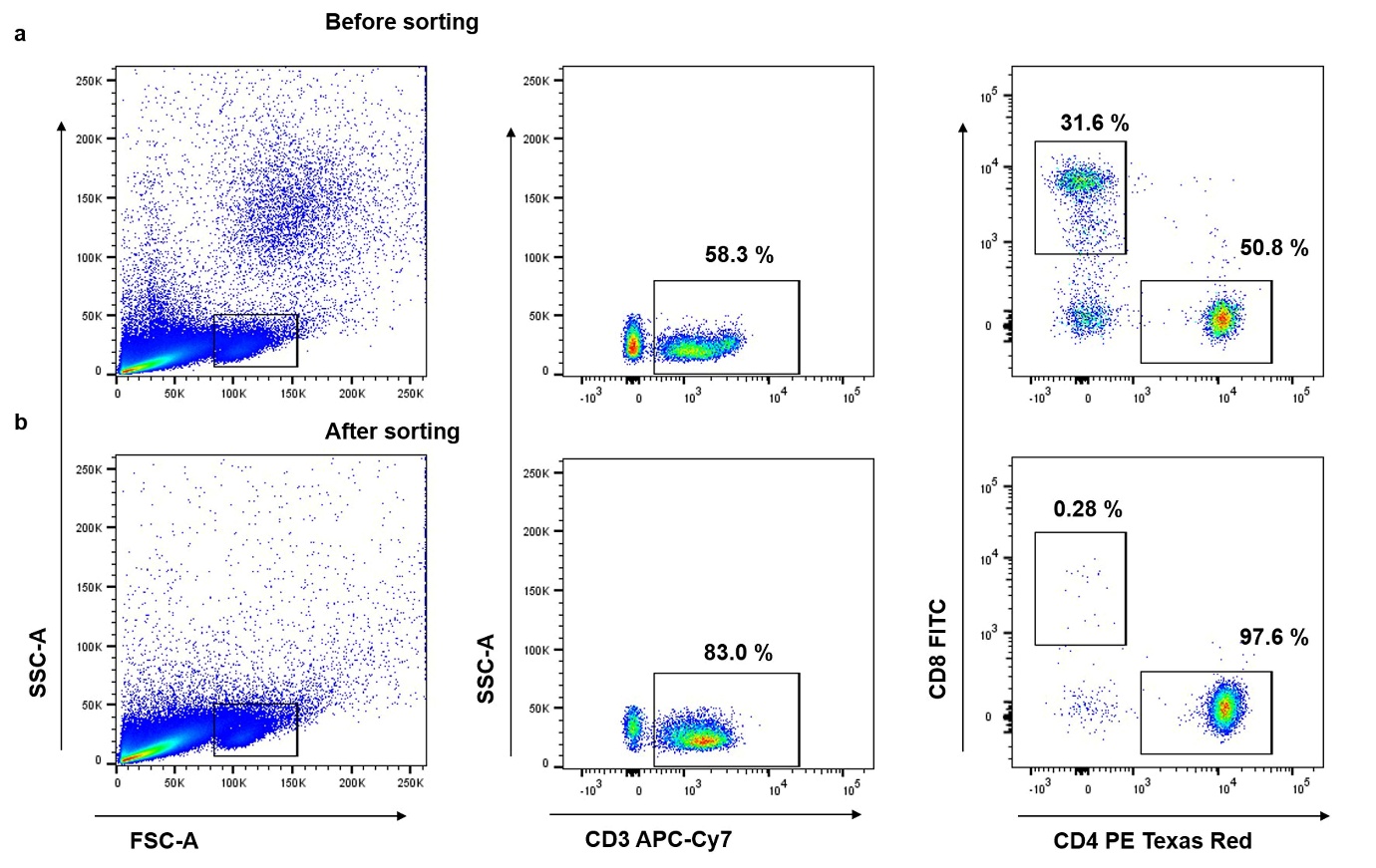

**Fig. S1**. **Enrichment of CD4⁺ Th cells from peripheral blood of perioperative breast cancer patients.** Representative flow cytometric analysis of CD3^+^CD4^+^CD8^-^ Th cells before (a) and after (b) sorting using MojoSort™ Human T Cell Isolation Kit, which enables the isolation of untouched CD4⁺ Th cells from peripheral blood.

**Validation of liquid phase administration of isoflurane**

To ascertain whether the observed apoptotic effects were specifically linked to isoflurane exposure rather than to nonspecific experimental conditions, we incorporated both vehicle and positive controls along with normal controls in our *in vitro* studies. Jurkat T cells treated with the vehicle control (VC; 0.1% DMSO) exhibited no significant induction of apoptosis compared to control (only Jurkat T cells without DMSO), thereby confirming that the solvent itself did not influence the observed effects (Fig. S2a, b). Conversely, hydrogen peroxide (H₂O₂) was employed as a positive control under the same experimental conditions to confirm the sensitivity of the apoptosis assays. Isoflurane significantly induced apoptosis of Jurkat T cells compared to control, vehicle control and propofol (Fig. S2a, b). These controls illustrate that the apoptotic response was specifically related to isoflurane exposure and not to the experimental setup or vehicle treatment.

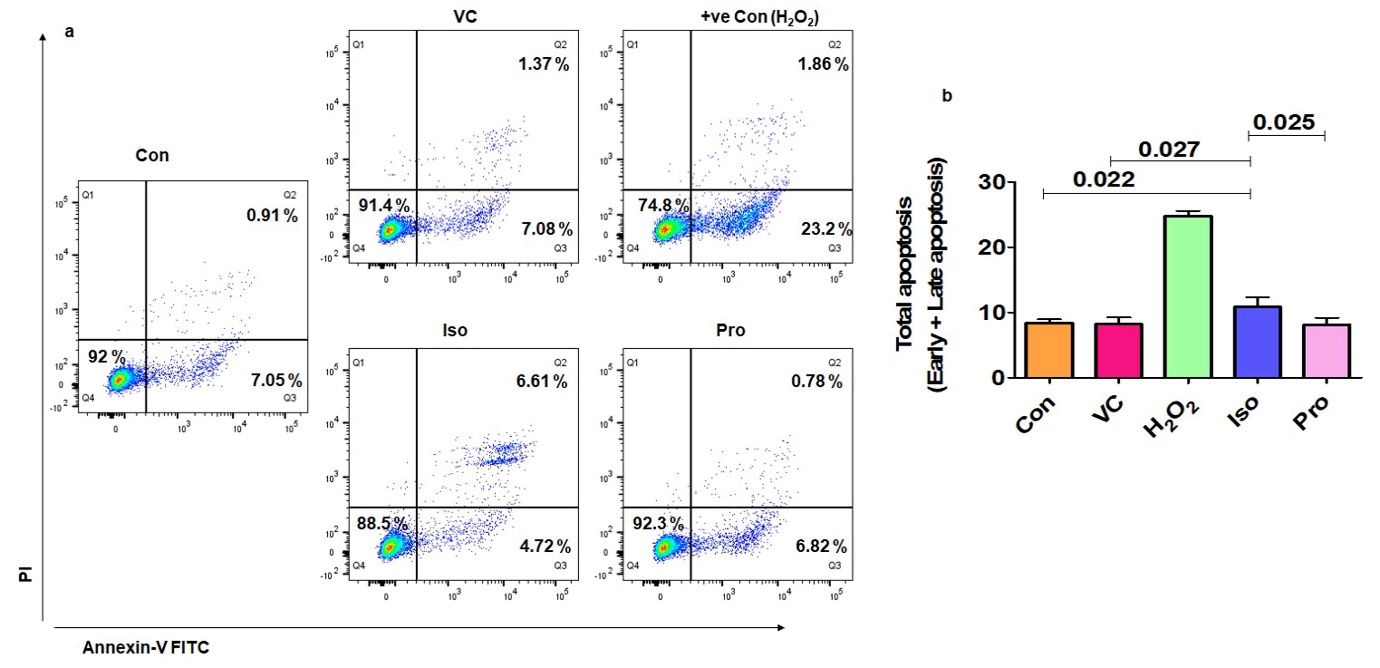

**Fig. S2:** Comparison of apoptosis between control, VC, +ve Con (H_2_O_2_), Iso and Pro in Jurkat T cells. Representative flow cytometric analysis of apoptosis (a) and comparative analysis on the effect of isoflurane/propofol on the total apoptosis of Jurkat T cells (b). [Con, Control; VC, Vehicle control; H_2_O_2_, Hydrogen peroxide; Iso, Isoflurane and Pro, Propofol]

***Table S1: List of RT-qPCR Primers***

| Gene | Primer (Human) | Sequence (5’- 3’) | Product Length | Accession Number | Annealing Temperature |
| --- | --- | --- | --- | --- | --- |
| *BAK1* | Forward | TGAGTACTTCACCAAGATTGCCA | 121 bps | NM_001188.4 | 59°C |
|  | Reverse | AGTCAGGCCATGCTGGTAGAC |  |  |  |
| *BAX* | Forward | CCCGAGAGGTCTTTTTCCGAG | 155 bps | NM_138761.4 | 52°C |
|  | Reverse | CCAGCCCATGATGGTTCTGAT |  |  |  |
| *BCL2* | Forward | GGTGGGGTCATGTGTGTGG | 89 bps | NM_000633.3 | 60°C |
|  | Reverse | CGGTTCAGGTACTCAGTCATCC |  |  |  |
| *CASP3* | Forward | GTAGATGGTTTGAGCCTGAG | 106 bps | NM_001354779.2 | 55°C |
|  | Reverse | CCAGTGCGTATGGAGAAATG |  |  |  |
| *CASP8* | Forward | CCTCCCTCAAGTTCCTGAGCCT | 209 bps | NM_001400666.1 | 57°C |
|  | Reverse | TTCCCTTTCCATCTCCTCCTTTCT |  |  |  |
| *FAS* | Forward | ATGCTGGGCATCTGGAC | 96 bps | NM_152871.4 | 60°C |
|  | Reverse | GGAGTTGATGTCAGTCACTT |  |  |  |
| *MCL1* | Forward | CCAAGAAAGCTGCATCGAACCAT | 151 bps | NM_001197320.2 | 55°C |
|  | Reverse | CAGCACATTCCTGATGCCACCT |  |  |  |
| *PARP1* | Forward | CCTGATCCCCCACGACTTT | 92 bps | NM_001618.4 | 56°C |
|  | Reverse | GCAGGTTGTCAAGCATTTC |  |  |  |
| *MAPK1* | Forward | CGTGTTGCAGATCCAGACCATGAT | 124 bps | NM_138957.3 | 61°C |
|  | Reverse | TGGACTTGGTGTAGCCTTTGGAA |  |  |  |
| *MAPK14* | Forward | AACCTGTCTCCAGTGGGCTCT | 71 bps | NM_139013.3 | 57°C |
|  | Reverse | CGTAACCCCGTTTTTGTGTCA |  |  |  |
| *MAPK9* | Forward | AGCCCAAGGGATTGTTTGTG | 123 bps | NM_001364607.2 | 56°C |
|  | Reverse | AGGACGAGTTCACGATAAGCTC |  |  |  |
| *GAPDH* | Forward | GTCTCCTCTGACTTCAACAGCG | 131 bps | NM_002046.7 | 55°C |
|  | Reverse | ACCACCCTGTTGCTGTAGCCAA |  |  |  |

[Abbreviations: *BAK1,* BCL2 antagonist/killer 1*; BAX*, BCL2 associated X*; BCL2,* B-cell lymphoma-2*; CASP3,* Caspase-3*; CASP8,* Caspase-8*; FAS*, Fas Cell Surface Death Receptor*;* GAPDH, Glyceraldehyde-3-phosphate dehydrogenase*; MAPK1*, Mitogen-activated protein kinase *1; MCL1,* MCL1 apoptosis regulator*; PARP1,* Poly (ADP-ribose) polymerase 1]

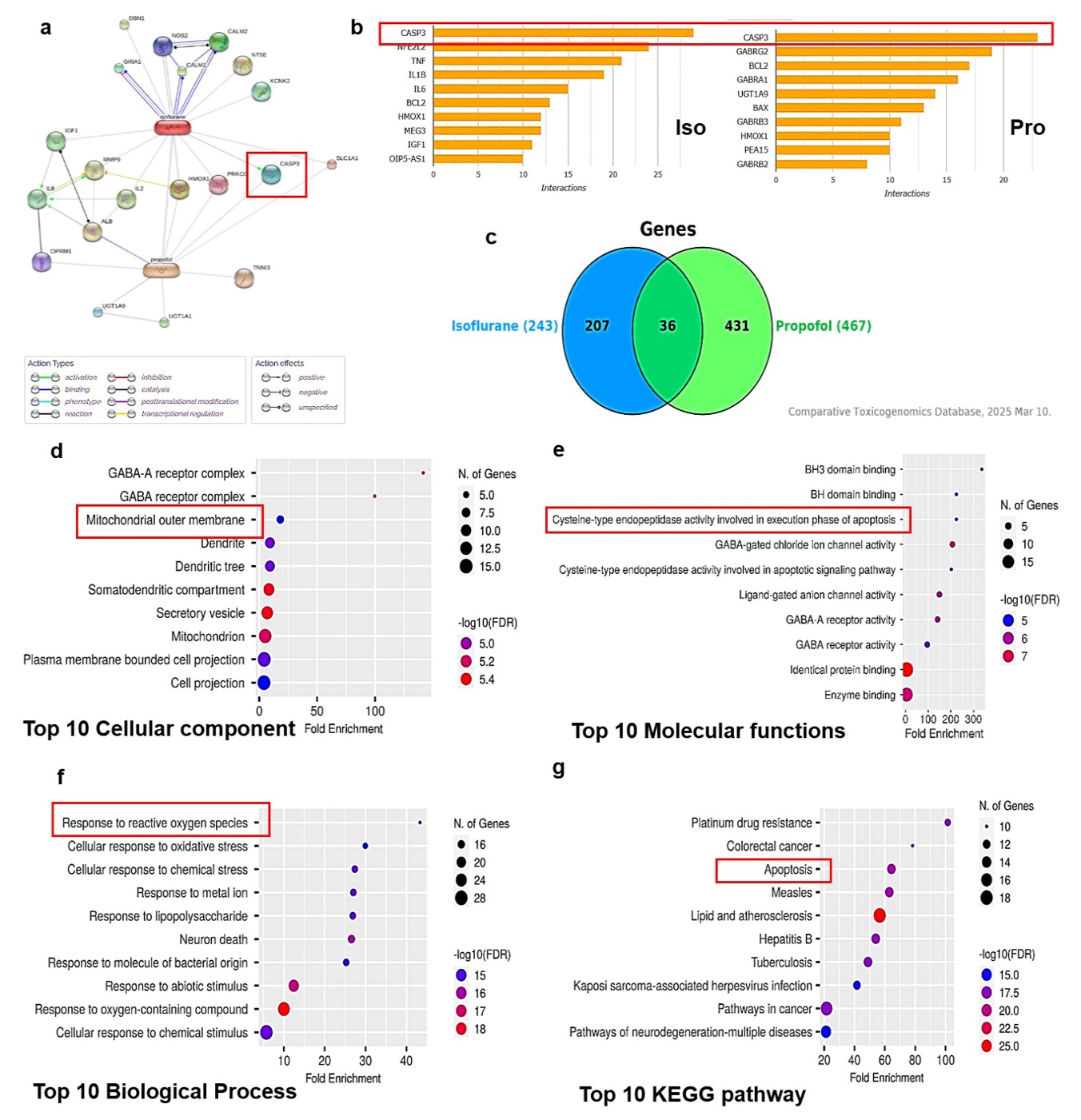

**Fig. S3:** **Bioinformatic identification of anesthetic agent-associated molecular targets and pathway enrichment analysis**. STITCH-based network analysis showing predicted chemical–protein interactions of isoflurane and propofol. Four common target proteins- HMOX1, PRKCG, CASP3, and SLC1A1 were identified (a); CTD analysis displaying the top-ranked genes interacting with isoflurane (left) and propofol (right), with *CASP3* emerging as the most strongly associated gene for both anesthetics (b); Venn diagram illustrating overlap between DEGs linked to isoflurane (N = 243) and propofol (N = 467), revealing 36 shared genes (c) from CTD analysis; Functional enrichment analysis of the 36 shared genes using ShinyGO v0.82. Top 10 enriched GO terms were ranked by fold enrichment for cellular component (d), molecular function (e), biological process (f) and KEGG pathway (g). [CASP3, Caspase-3; CTD, Comparative Toxicogenomics Database; HMOX1, Heme oxygenase 1; Iso, Isoflurane; KEGG, Kyoto Encyclopedia of Genes and Genomes; PRKCG, Protein kinase C gamma; Pro, Propofol; SLC1A1, Solute carrier family 1 member 1]

***Table S2: Common Interacting Proteins with both Isoflurane and Propofol***

| Protein | Drug | Prediction for specific actions | Combined Score |
| --- | --- | --- | --- |
| HMOX1 | Iso | None | 0.982 |
|  | Pro |  | 0.818 |
| PRKCG | Iso |  | 0.843 |
|  | Pro |  | 0.817 |
| CASP3 | Iso | Activation (0.700) | 0.956 |
|  | Pro | None | 0.826 |
| SLC1A1 | Iso |  | 0.843 |
|  | Pro |  | 0.800 |

[Abbreviations: HMOX1, heme oxygenase 1; PRKCG, protein kinase C gamma; CASP3, caspase-3; SLC1A1, solute carrier family 1 member 1]

***Table S3: Clinico-Pathological Features of Treatment Naive Breast Cancer Patients Enrolled for Surgical Resection in Two Different Anesthestic Groups- Isoflurane and Propofol***

| Characters | Character subtype | Anesthetic agent | | *p* value |
| --- | --- | --- | --- | --- |
|  |  | **Isoflurane**  **(N, 15)** | **Propofol**  **(N, 15)** |  |
| Age (yr) | | 57.66 ± 13.17 | 57.13 ± 11.73 | 0.908 |
| Height (cm) | | 148.80 ± 4.29 | 147.26 ± 9.85 | 0.585 |
| Weight (kg) | | 52.40 ± 7.89 | 52.00 ± 6.95 | 0.884 |
| ASA (%) | ASA I | 53.33 | 33.33 | 0.157 |
|  | ASA II | 46.66 | 66.66 |  |
| Histopathology (%) | Invasive carcinoma of NOS type | 26.66 | 53.33 | 0.199 |
|  | Invasive/ Infiltrating ductal carcinoma | 66.66 | 33.33 |  |
|  | Ductal carcinoma in situ | 6.66 | 6.66 |  |
|  | Mucinous carcinoma |  | 6.66 |  |
| Molecular subtypes (%) | ER^+^PR^+^HER2^-^ | 33.33 | 53.33 | 0.238 |
|  | ER^-^PR^-^HER2^+^ | 13.33 |  |  |
|  | ER^-^PR^-^HER2^-^ | 33.33 | 6.66 |  |
|  | ER^+^PR^+^HER2^+^ | 6.66 | 13.33 |  |
|  | ER^+^PR^-^HER2^-^ | 13.33 | 20.00 |  |
|  | ER^+^PR^-^HER2^+^ |  | 6.66 |  |
| Type of surgery (%) | MRM | 66.66 | 53.33 | 0.157 |
|  | BCS | 33.33 | 46.66 |  |
| Duration of surgery (min) | | 88.66 ± 26.28 | 91.33 ± 36.91 | 0.821 |
| Duration of anesthesia (min) | | 108.66 ± 27.80 | 116.66 ± 31.49 | 0.467 |
| Pain score (Number) | | 1.66 ± 0.62 | 1.80 ± 0.56 | 0.541 |
| Grade (%) | Grade I | 6.66 | 26.66 | 0.083 |
|  | Grade II | 86.66 | 46.66 |  |
|  | Grade III | 6.66 | 26.66 |  |
| Stage (%) | Stage II | 80.00 | 66.66 | 0.157 |
|  | Stage III | 20.00 | 33.33 |  |

**Data represented as mean ± SD or percentage (%); Student’s t-test and Chi-square test has been performed (as applicable to specific parameter) to compare the clinicopathological characteristics.*

***Table S4: Linear Mixed Effects Regression Showing Effect of Isoflurane on Apoptosis Markers***

| Parameters | β-estimate | 95% CI | p-value |
| --- | --- | --- | --- |
| *Apoptosis* | | | |
| Isoflurane | -4.375 | -13.422 to 4.671 | 0.3472 |
| Timepoints Intra | -4.217 | -11.58 to 3.147 | 0.2665 |
| Timepoints Post | -6.845 | -14.209 to 0.518 | 0.0738 |
| Age | -0.113 | -0.385 to 0.16 | 0.4277 |
| BMI | 1.042 | -0.039 to 2.122 | 0.072 |
| Stage 2 | 0.71 | -6.717 to 8.136 | 0.8532 |
| Duration of anesthesia | -0.01 | -0.129 to 0.108 | 0.8635 |
| Surgery type 2 | -3.535 | -10.371 to 3.3 | 0.3217 |
| ASA2 | -1.762 | -8.868 to 5.344 | 0.6318 |
| Isoflurane: Timepoints Intra | **15.934** | **5.52 to 26.348** | **0.004** |
| Isoflurane: Timepoints Post | 7.709 | -2.705 to 18.122 | 0.1524 |
| *JC-1* | | | |
| Isoflurane | 0.029 | -0.548 to 0.605 | 0.9222 |
| Timepoints Intra | 0.093 | -0.377 to 0.564 | 0.6988 |
| Timepoints Post | -0.197 | -0.668 to 0.273 | 0.4144 |
| Age | 0.006 | -0.011 to 0.024 | 0.475 |
| BMI | 0.029 | -0.039 to 0.098 | 0.4114 |
| Stage 2 | -0.208 | -0.68 to 0.264 | 0.3976 |
| Duration of anesthesia | -0.001 | -0.008 to 0.007 | 0.8359 |
| Surgery type 2 | 0.288 | -0.146 to 0.723 | 0.2073 |
| ASA2 | -0.176 | -0.628 to 0.276 | 0.453 |
| Isoflurane: Timepoints Intra | **-1.735** | **-2.401 to -1.07** | **0.000** |
| Isoflurane: Timepoints Post | -0.419 | -1.084 to 0.247 | 0.2225 |
| *TUNEL* | | | |
| Isoflurane | -4.502 | -61.038 to 52.034 | 0.8766 |
| Timepoints Intra | 3.267 | -37.713 to 44.247 | 0.8764 |
| Timepoints Post | 16.8 | -24.18 to 57.78 | 0.4251 |
| Age | 0.301 | -1.539 to 2.141 | 0.7514 |
| BMI | 3.41 | -3.873 to 10.694 | 0.3687 |
| Stage 2 | 3.687 | -46.375 to 53.75 | 0.8865 |
| Duration of anesthesia | -0.177 | -0.974 to 0.619 | 0.6665 |
| Surgery type 2 | -7.44 | -53.518 to 38.639 | 0.7547 |
| ASA2 | 4.314 | -43.59 to 52.217 | 0.8615 |
| Isoflurane: Timepoints Intra | **160.467** | **102.512 to 218.421** | **0.000** |
| Isoflurane: Timepoints Post | 35.6 | -22.355 to 93.555 | 0.2337 |
| *ROS* | | | |
| Isoflurane | 110.163 | -56.239 to 276.565 | 0.2001 |
| Timepoints Intra | -79.867 | -210.242 to 50.508 | 0.2349 |
| Timepoints Post | -86.267 | -216.642 to 44.108 | 0.2 |
| Age | -0.578 | -5.741 to 4.585 | 0.8284 |
| BMI | 2.225 | -18.218 to 22.667 | 0.8331 |
| Stage 2 | 37.133 | -103.371 to 177.637 | 0.6096 |
| Duration of anesthesia | -1.179 | -3.415 to 1.056 | 0.3123 |
| Surgery type 2 | 9.182 | -120.141 to 138.505 | 0.8906 |
| ASA2 | -5.602 | -140.047 to 128.842 | 0.9356 |
| Isoflurane: Timepoints Intra | **409.933** | **225.555 to 594.312** | **0.0001** |
| Isoflurane: Timepoints Post | 126.733 | -57.645 to 311.112 | 0.1833 |
| *Caspase-3/7 activity* | | | |
| Isoflurane | -276.33 | -628.182 to 75.522 | 0.1326 |
| Timepoints Intra | -27.933 | -232.387 to 176.52 | 0.7899 |
| Timepoints Post | **-530.733** | **-735.187 to -326.28** | **0.000** |
| Age | -6.86 | -19.363 to 5.644 | 0.2939 |
| BMI | -13.485 | -62.99 to 36.021 | 0.5988 |
| Stage 2 | 50.063 | -290.196 to 390.322 | 0.7758 |
| Duration of anesthesia | -0.856 | -6.269 to 4.556 | 0.7594 |
| Surgery type 2 | **404.673** | **91.491 to 717.855** | **0.019** |
| ASA2 | 19.487 | -306.096 to 345.071 | 0.9077 |
| Isoflurane: Timepoints Intra | **1335.933** | **1046.792 to 1625.075** | **0.000** |
| Isoflurane: Timepoints Post | **1514.933** | **1225.792 to 1804.075** | **0.000** |
| *Bcl2 (Protein expression)* | | | |
| Isoflurane | 0.013 | -0.168 to 0.193 | 0.8917 |
| Timepoints Intra | **0.18** | **0.018 to 0.342** | **0.034** |
| Timepoints Post | **0.177** | **0.015 to 0.34** | **0.0366** |
| Age | 0.003 | -0.002 to 0.008 | 0.1926 |
| BMI | -0.001 | -0.02 to 0.019 | 0.9448 |
| Stage 2 | 0.129 | -0.006 to 0.264 | 0.0744 |
| Duration of anesthesia | -0.001 | -0.003 to 0.002 | 0.615 |
| Surgery type 2 | 0.094 | -0.03 to 0.219 | 0.1496 |
| ASA2 | -0.09 | -0.219 to 0.039 | 0.1863 |
| Isoflurane: Timepoints Intra | -0.145 | -0.375 to 0.084 | 0.2198 |
| Isoflurane: Timepoints Post | -0.201 | -0.431 to 0.028 | 0.0911 |
| *Bax (Protein expression)* | | | |
| Isoflurane | -0.062 | -0.212 to 0.088 | 0.4229 |
| Timepoints Intra | -0.071 | -0.179 to 0.037 | 0.2009 |
| Timepoints Post | -0.108 | -0.216 to 0.00 | 0.055 |
| Age | 0.00 | -0.005 to 0.005 | 0.9673 |
| BMI | 0.004 | -0.016 to 0.023 | 0.7182 |
| Stage 2 | -0.097 | -0.23 to 0.036 | 0.167 |
| Duration of anesthesia | -0.001 | -0.003 to 0.001 | 0.5031 |
| Surgery type 2 | 0.121 | -0.001 to 0.243 | 0.0655 |
| ASA2 | -0.056 | -0.184 to 0.071 | 0.3936 |
| Isoflurane: Timepoints Intra | **0.384** | **0.231 to 0.537** | **0.000** |
| Isoflurane: Timepoints Post | **0.175** | **0.023 to 0.328** | **0.0284** |
| *Mcl-1 (Protein expression)* | | | |
| Isoflurane | -0.251 | -0.982 to 0.481 | 0.5047 |
| Timepoints Intra | 0.327 | -0.265 to 0.92 | 0.2836 |
| Timepoints Post | -0.117 | -0.709 to 0.476 | 0.7011 |
| Age | -0.018 | -0.04 to 0.004 | 0.1232 |
| BMI | -0.087 | -0.174 to 0.001 | 0.0657 |
| Stage 2 | -0.214 | -0.817 to 0.388 | 0.4929 |
| Duration of anesthesia | 0.003 | -0.007 to 0.012 | 0.6055 |
| Surgery type 2 | **0.685** | **0.13 to 1.239** | **0.0242** |
| ASA2 | -0.118 | -0.695 to 0.458 | 0.6917 |
| Isoflurane: Timepoints Intra | -0.412 | -1.25 to 0.426 | 0.3394 |
| Isoflurane: Timepoints Post | 0.085 | -0.753 to 0.923 | 0.8438 |
| *cleaved Caspase-3 (Protein expression)* | | | |
| Isoflurane | -0.053 | -0.198 to 0.091 | 0.473 |
| Timepoints Intra | -0.093 | -0.233 to 0.046 | 0.1941 |
| Timepoints Post | -0.086 | -0.225 to 0.053 | 0.231 |
| Age | -0.002 | -0.006 to 0.001 | 0.2434 |
| BMI | -0.001 | -0.015 to 0.014 | 0.9218 |
| Stage 2 | -0.057 | -0.155 to 0.041 | 0.2699 |
| Duration of anesthesia | -0.001 | -0.002 to 0.001 | 0.3682 |
| Surgery type 2 | 0.054 | -0.036 to 0.144 | 0.252 |
| ASA2 | -0.016 | -0.11 to 0.078 | 0.7399 |
| Isoflurane: Timepoints Intra | **0.347** | **0.15 to 0.544** | **0.001** |
| Isoflurane: Timepoints Post | 0.016 | -0.181 to 0.213 | 0.874 |
| *cleaved PARP1 (Protein expression)* | | | |
| Isoflurane | -0.018 | -0.546 to 0.51 | 0.9476 |
| Timepoints Intra | 0.214 | -0.18 to 0.608 | 0.2914 |
| Timepoints Post | -0.147 | -0.541 to 0.246 | 0.4664 |
| Age | 0 | -0.017 to 0.017 | 0.9896 |
| BMI | -0.012 | -0.079 to 0.055 | 0.7349 |
| Stage 2 | 0.103 | -0.357 to 0.563 | 0.6639 |
| Duration of anesthesia | 0.006 | -0.001 to 0.013 | 0.1299 |
| Surgery type 2 | **0.556** | **0.132 to 0.979** | **0.0174** |
| ASA2 | -0.056 | -0.496 to 0.384 | 0.8061 |
| Isoflurane: Timepoints Intra | **0.608** | **0.051 to 1.165** | **0.0367** |
| Isoflurane: Timepoints Post | 0.143 | -0.414 to 0.7 | 0.6175 |

*[Propofol was considered the reference category for the anesthetic group variable, and estimates for the isoflurane group were interpreted relative to propofol. Multicollinearity among the independent variables was assessed using the variance inflation factor (VIF) derived from a linear regression model including all covariates. The VIF values for all predictors were close to 1 (range: 1.00–1.19), indicating no significant correlation among the explanatory variables (age, BMI, ASA, duration of anesthesia, type of surgery and stage). As all VIF values were well below the commonly accepted threshold of 5, there was no evidence of multicollinearity.]*

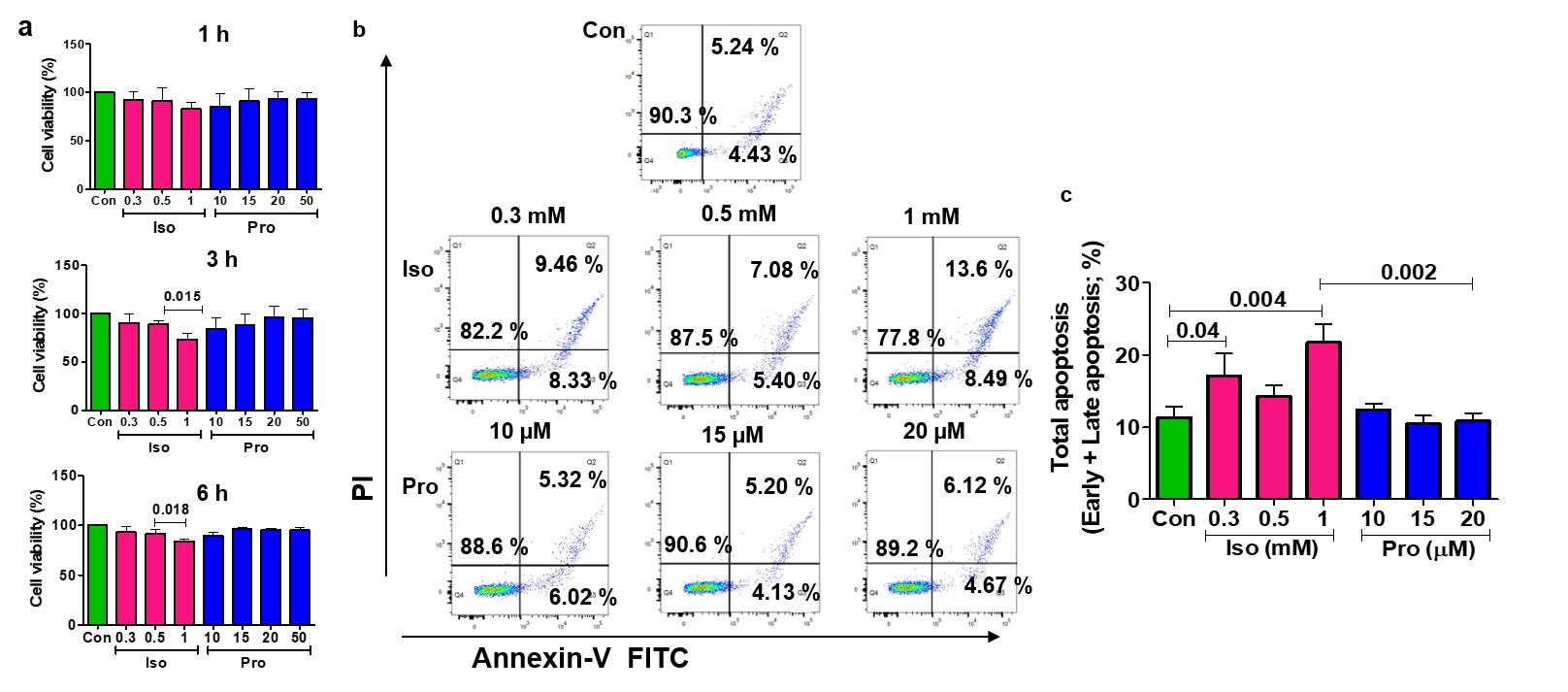

**Fig. S4:** **Concentration and time dependent studies of isoflurane and propofol on cell viability and apoptosis of Jurkat T cells**. Effect of isoflurane (0.3-1 mM) and propofol (10-50 µM) on cell viability of Jurkat T cells at 1, 3 and 6 h (a); Flow cytometric representation of the effect of isoflurane (0.3-1 mM) and propofol (10-20 µM) on apoptosis of Jurkat T cells at 3 h (b) and comparative analysis on the effect of isoflurane/propofol on the total apoptosis of Jurkat T cells (c) at different concentrations. [The graphs were plotted based on the mean± SD; Con, Control; Iso, Isoflurane; Pro, Propofol].
